# Serological evidence of high pathogenicity virus infection in *Eidolon helvum* fruit bats in Nigeria

**DOI:** 10.1101/2023.06.14.544916

**Authors:** Diego Cantoni, Martin Mayora-Neto, Bethany Auld, Kelly Da Costa, Joanne Del Rosario, Veronica O. Ameh, Claude T. Sabeta, Mariliza Derveni, Arran Hamlet, Edward Wright, Simon Scott, Efstathios S. Giotis, Ashley Banyard, Nigel Temperton

**Author notes:** These senior authors were co-principle investigators (Efstathios S. Giotis, Ashley Banyard, Nigel Temperton).

## Abstract

The *Eidolon helvum* fruit bat is the most widely distributed fruit bat in Africa and is known to be a reservoir for several pathogenic viruses that can cause disease in humans. To assess the risk of zoonotic spillover, we conducted a serological survey of 304 serum samples from *E. helvum* bats that were captured for human consumption in Makurdi, Nigeria. Using pseudotyped viruses, we screened the samples for neutralising antibodies against viruses from the *Coronaviridae, Filoviridae, Orthomyxoviridae* and *Paramyxoviridae* families. We report the presence of neutralising antibodies against henipavirus lineage GH-M74a virus (odds ratio 6.23; p<0.001), Nipah virus (odds ratio 4.04; p=0.00031), bat influenza H17N10 virus (odds ratio 7.25; p<0.001) and no significant association with Ebola virus (odds ratio 0.56; p=0.375) in the bat cohort. The data suggest a potential risk of zoonotic spillover including the possible circulation of highly pathogenic viruses in *E. helvum* populations. These findings highlight the importance of maintaining sero-surveillance of *E. helvum* to monitor changes in virus prevalence and distribution over time and across different geographic locations.

**Article summary line:** The detection of neutralizing antibodies against henipavirus GH-M74a virus, Nipah virus, and H17N10 virus in *Eidolon helvum* bat sera from Nigeria using pseudotyped viruses suggests a potential risk of zoonotic spillover.

## Introduction

Throughout the course of human history, viral zoonotic spillover events have been sporadic yet catastrophic, resulting in several highly lethal pandemics. With approximately 60-75% of all human infectious diseases arising from zoonotic transmission (*1*), it is crucial to remain vigilant in identifying and monitoring potential sources of zoonotic spillover, such as wildlife reservoirs, in order to prevent future outbreaks. Over the past four decades, bats have been identified as a significant source of zoonotic events that have sparked major outbreaks of viruses with considerable implications for human health. As the second most diverse mammalian order, bats have been linked to the transmission of a range of viruses, including coronaviruses, filoviruses, lyssaviruses, and henipaviruses, among others (*1,2*). With over 12,000 bat-derived virus sequences spanning 30 viral families having relevance to both veterinary and medical sectors, the need for surveillance of bat populations is critical to assess and mitigate the risk of potential zoonotic spillover events.

*Eidolon helvum*, the straw-coloured fruit bat, is one of the most widely distributed fruit bats in Africa (Figure 1A). It has extensive migratory patterns of over 2000 kilometres and is hypothesised to migrate based on availability of fruits to increase reproductive success (*3*). It is often hunted for either its bushmeat (*4,5*) or for pest control (*6*). Due to the overlap of territories shared between *E. helvum* and humans (Figure 1B), there is ample opportunity for zoonotic spillover. According to the DBatVir database, more than 900 viruses have been sequenced from *E. helvum* (Figure 1C). Furthermore, the presence of neutralising antibodies have been detected against multiple viruses in serum samples of *E. helvum* (*7*).

**Figure 1.**
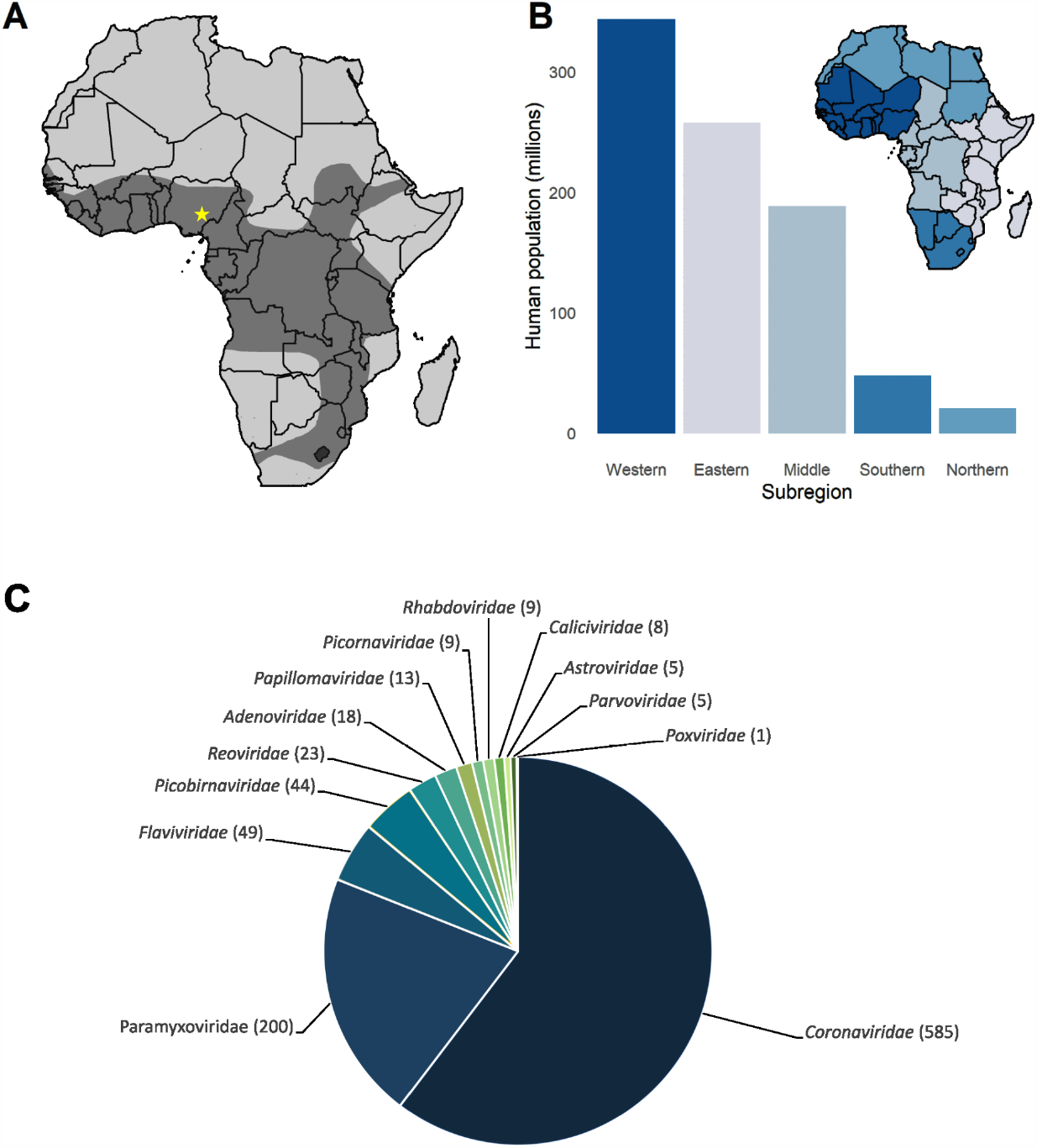
Profile of Straw-coloured *E. helvum* fruit bat. Species distribution within Africa coloured in dark grey, with yellow star denoting location in Nigeria from which the samples were obtained (A). Human population density within each subregion of Africa (B). Viruses reported in *E. helvum* samples in DBatVir database (http://www.mgc.ac.cn/cgi-bin/DBatVir/).

Serological evidence of neutralizing antibodies in bat sera is not definitive proof of active virus infection in bats, but it does suggest that bats have been exposed to the virus and have mounted an immune response. Screening for neutralising antibodies in bats using highly pathogenic viruses requires use of high containment facilities, which can be circumvented by using pseudotyped viruses (PVs) in neutralization assays. The use of PVs in neutralization assays is considered safe for handling under biosafety level 2 conditions. Pseudotyped virus neutralization assays (PVNA) have gained widespread usage for detecting neutralizing antibodies due to their high sensitivity and robust correlation with live virus neutralization assays (*8–10*). In this study, we screened 304 serum samples from *E. helvum* bats that were captured for human consumption in Nigeria using PVs expressing the viral glycoproteins of several viruses known to pose high public health risks (Table 1). Due to limited volumes of sera, we prioritised the order of screening based on the viruses’ potential risk to animal and human health (*11–13*). Our findings revealed the presence of multiple virus-specific neutralizing antibodies, suggesting the circulation of highly pathogenic viruses with pandemic potential in colonies of *E. helvum* in Nigeria.

**Table 1.**
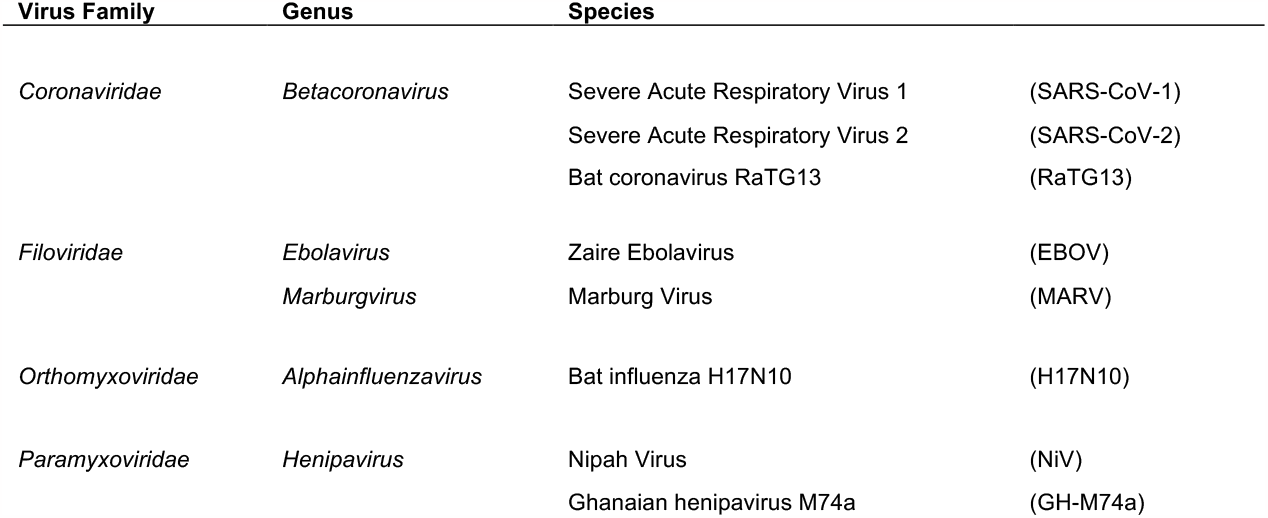
List of viruses pseudotyped for screening *E. helvum* samples in this study.

## Materials and Methods

### Bat sera collection

All sera were collected from terminally bled straw-coloured fruit bats (*Eidolon helvum*) that were captured for human consumption in Makurdi, Benue State Nigeria (7°44’25.7’’N 8°31’52.8’’E). The bats in Makurdi were collected from roosts in trees in and around the Benue State Government House and on trees in private residences close to the government house. Bats were also sampled from roosts on trees in private residences where permissions were gained. Sampling was done for two consecutive seasons (November 2017 – March 2018 and November 2018 – March 2019). Blood samples were processed for sera and stored at -20°C until required.

### Pseudotype virus production

Lentiviral (HIV) based pseudotypes used for this study were generated as described in detail here (*14,15*). Briefly, 1.0 μg of p8.91 plasmid encoding the HIV gag-pol was mixed with 1.5 μg of pCSFLW reporter gene and 1 ug the surface viral glycoproteins as required (Table 2). After mixing the plasmids in 200 μL of Opti-MEM (ThermoFisher, Woolwich, UK), Fugene HD (Promega, Southampton, UK) transfection reagent was added at a 1:3 (plasmid:Fugene HD) ratio and incubated for 15 minutes prior to adding the transfection complexes to HEK293T/17 cells in T-75 cell culture flasks. Harvesting of pseudotypes were carried out at 48- and 72-hours post transfection, whereby culture media was removed from the flasks and filtered through a 0.45 μM cellulose acetate filter (Corning, Deeside, UK). Samples were aliquoted and frozen at -80°C for long term storage prior to use.

**Table 2.** Plasmids used to generate pseudotyped viruses. HEK293T Cells were transfected with Angiotensin-converting enzyme 2 (ACE2) and Transmembrane protease, serine 2 (TMPRSS2). CHO: Chinese hamster ovary cells. MDCK: Madin-Darby canine kidney cells. For this study we used the G and F genes from a Bangladeshi NiV isolate (NiVb)

| Viral Glycoprotein | Accession | Vector | Target Cells |
| --- | --- | --- | --- |
| *Coronaviridae* |  |  |  |
| SARS-CoV S (Tor2) | NC_004718.3 | pcDNA 3.1+ | HEK293T ACE2+TMPRSS2 |
| SARS-CoV-2 S (Wuhu1) | NC_045512.2 | pcDNA 3.1+ | HEK293T ACE2+TMPRSS2 |
| RaTG13 S | QHR63300.2 | pcDNA 3.1+ | HEK293T ACE2+TMPRSS2 |
| *Filoviridae* |  |  |  |
| EBOV Mayinga 76 GP | EU224440 | pCAGGS | CHO |
| MARV Angola 05 GP | DQ447660 | pCAGGS | CHO |
| *Orthomyxoviridae* |  |  |  |
| IAV-like H17N10 H17 | AFC35438.1 | pl.18 | MDCK II |
| *Paramyxoviridae* |  |  |  |
| NiVb G and F | JN808864.1 | pCAGGS | BHK |
| GH-M74A G and F | HQ660129 | pCAGGS | BHK |

### Pseudotype virus titration

Pseudotyped viruses were titrated by serially diluting filtered pseudotypes with fresh DMEM starting from a 1:2 to a final 1:512 dilution using white 96-well flat bottom plates (ThermoFisher, Woolwich, UK). Then appropriate target cells were added (Table 1) and plates were returned to the incubator. After 48 hours, cells were lysed with Bright-Glo reagent (Promega, Southampton, UK), and luciferase expression levels were assessed using a GloMax plate reader (Promega, Southampton, UK).

### Sera screen and neutralisations

Sera were initially screened at single point dilution (1:100, except NiV and GH-M74a at 1:50). Each plate had wells containing PV only to determine maximum pseudotype entry. Samples where a 1 log decrease in RLU compared to the no-serum virus-only control was observed were then selected for cytotoxicity assay using Cell Titre Glo kit (Promega, Southampton, UK) and light microscopy to verify viability of cells prior to undertaking PVNA (data not shown). Neutralisation assays were carried out by mixing bat sera and cell culture media at a starting input of 1:40 ratio (sera: cell culture media) and diluted either 4-fold (GH-M74a and NiV) to 1:5000 or 8-fold (all other viruses) to 1:5120. PVs were added to the plates at a minimum RLU titre of 10^5^/mL, incubated for 1 hour at 37°C, followed by addition of target cells at a density of 10^4^ cells per well, except for NiV and GH-M74a where a cell density of 20^4^ cells per well. Plates were incubated at 37°C and 5% CO2 for 48 hours prior to lysis using Bright-Glo reagent according to manufacturer’s protocol and monitoring of luciferase expression using a GloMax plate reader. Regression curves were fitted using GraphPad Prism 8 software (San Diego, CA, USA) as described previously (*16*).

### Statistical analysis

Data were analyzed using STATA version 16 (StataCorp, College Station, TX, USA). Descriptive statistics were used to summarize the distribution of the variables. Bivariate logistic regression was used to determine the association between the presence or absence of neutralizing antibodies against each virus. Multivariable logistic regression was used to adjust for confounding factors. The data were coded as binary variables (1 = positive, 0 = negative). Logistic regression was used to determine the association between the presence or absence of neutralizing antibodies against each virus and the dependent variable was the presence or absence of neutralizing antibodies against each virus. Crude and adjusted odds ratios (OR) were estimated with 95% confidence intervals (CI) to measure the strength of association between each variable and the presence or absence of neutralizing antibodies.

## Results

Our PVNA screening has revealed the presence of neutralising antibodies against several of the pseudotyped viruses tested (Figure 2A and 2B, Table 3). However, no neutralization was observed against the *Coronaviridae* members SARS-CoV-2, SARS-CoV, and the bat coronavirus RaTG13. Although neutralizing antibodies against EBOV PVs were detected in a single sample (n=1/278), no neutralization was observed against MARV. We also found several samples positive for neutralizing antibodies against influenza A virus H17N10 PVs (n=26/304), indicating the presence of positive or cross-reactive H17 neutralizing antibodies. Additionally, positive samples were detected for Nipah virus (NiV) (n=16/54) and GH-M74a (n=36/54) from the *Paramyxoviridae* PVs.

**Table 3.** Results of pMN assays with each PV. *Note: sample sizes decreased due to limited sera volumes*.

| Viruses | Positive Samples | Negative | Percentage |
| --- | --- | --- | --- |
| *Coronaviridae* |  |  |  |
| SARS-CoV-1 | 0 | 304 | 0.00% |
| SARS-CoV-2 | 0 | 290 | 0.00% |
| RaTG13 | 0 | 290 | 0.00% |
| *Filoviridae* |  |  |  |
| EBOV | 1 | 278 | 0.36% |
| MARV | 0 | 279 | 0.00% |
| *Orthomyxoviridae* |  |  |  |
| H17N10 | 26 | 304 | 8.55% |
| *Paramyxoviridae* |  |  |  |
| NiV | 16 | 54 | 29.63% |
| Gh-M74a | 36 | 54 | 66.67% |

**Figure 2.**
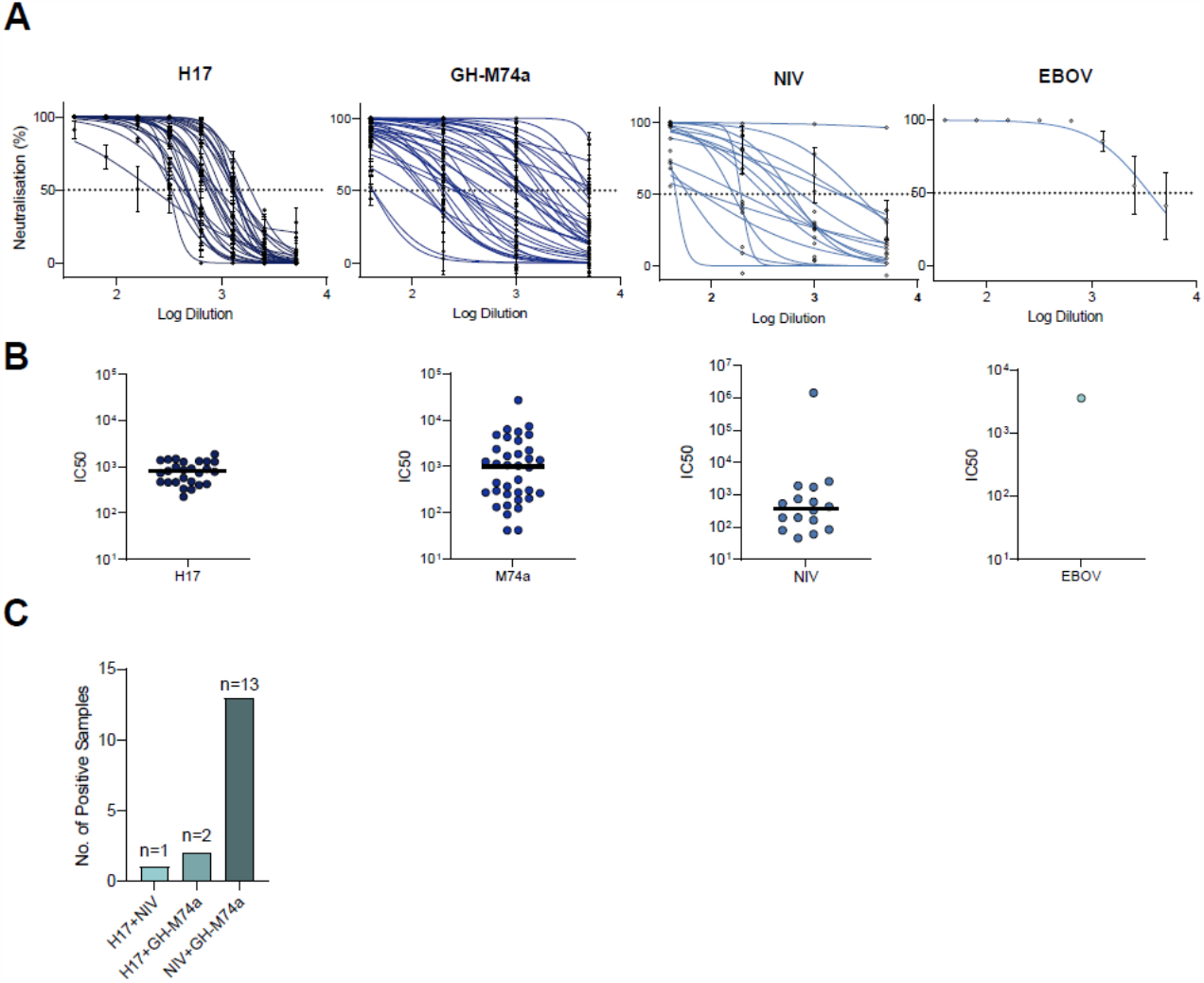
Positive detection of neutralising antibodies in *E. helvum* by pMN assay. Neutralisation curves derived from pMN assays against positive samples that were selected from the initial screening (A). IC50s calculated by pMN assays against each PV (B). Number of samples positive for neutralising antibodies against multiple viruses (C).

Our logistic regression analysis shows that the presence of neutralizing antibodies is significantly associated with virus type, as indicated by the p-values for the virus coefficients. Specifically, the odds of having neutralizing antibodies are significantly higher for H17 (odds ratio = 7.25, p < 0.001), NiV (odds ratio = 4.04, p = 0.00031), and GH-M74a (odds ratio = 6.23, p < 0.001) compared to the reference category of SARS-CoV-2. EBOV did not show a significant association with neutralizing antibody status (odds ratio = 0.56, p = 0.375).

We further analyzed the data to investigate the presence of samples positive for neutralizing antibodies against multiple viruses (Figure 2C). Our analysis revealed one sample positive for only H17, two samples positive for H17 and GH-M74a, and 13 samples positive for both NiV and GH-M74a. These findings suggest that multiple viruses may have circulated within the same bat or that antibodies to one of the three viruses may cross-neutralise against multiple viruses.

## Discussion

The study investigated the seroprevalence of neutralizing antibodies against several highly pathogenic viruses in *E. helvum* bats in Nigeria. We report a high level of seroprevalence against GH-M74a (66.7% of samples) and NiV (29.6%). This suggests that these viruses may be prevalent in the bat population, which is concerning given the high mortality rates (50-100%) associated with NiV infection in humans (*17*). Henipavirus antibodies have been frequently detected with high seroprevalence rates in *E. helvum* (18–20). We considered that the antibodies generated by GH-M74a infection are also cross-neutralizing the NiV viruses as the two viruses share a key conserved region in their attachment protein (*18*). Nonetheless, the detection of three unique samples that neutralized only NiV and not GH-M74a PV implies that NiV may indeed have been in circulation within this bat population. Currently no vaccines that can prevent nor treat NiV infection and disease have been licenced, though experimental mRNA-based vaccines are currently in development for NiV and are under phase 1 clinical trials. The study found a high level of detection of the GH-M74a virus in *E. helvum* bats, indicating that it may be circulating widely in the population. This suggests a risk of zoonotic spillover, potentially through secondary hosts such as pigs and horses, even if not directly to humans.

We also observed seropositivity towards H17N10, a bat Influenza A-like virus that contains unconventional HA and NA proteins (*21*). The evolutionary distinct H17N10 along with H18N11 virus (not screened for in this study) were originally recovered from asymptomatic fruit bats of the Neotropic bat family *Phyllostomidae* (*Sturnira lilium* and *Artibeus planirostris*) in several countries of Central and South America (*21–23*). These viruses have attracted considerable attention following reports that their entry in host cells is mediated by the trans-species conserved MHC-II proteins, suggesting zoonotic potential (*24,25*). So far, bats in Central and S. America, but not in Central Europe (*26*), have been found seropositive for H17N10 and H18N11, and to our knowledge this is the first report of H17-seropositive samples in non-Neotropical bat species. Considering the lack of a validated serological assay to screen *E. helvum* bat sera specifically for H17N10, we cannot exclude serological cross-reactivity with heterologous H17-like antigens. Nonetheless, the potential presence of undiscovered H17 or H17-like IAV species in *E. helvum*, such as the ones described in *Phylostomidae* is tantalising, cannot be ruled out and should be further investigated by unbiased approaches *i*.*e*., metagenomics.

Although we only detected a single sample positive for neutralising antibodies against EBOV, similar studies have also reported very low seroprevalence rates in *E. helvum;* 1 out of 262 samples from Ghana (*27*), and 19 out of 748 samples from Zambia (*28*). On the other hand, a report detected much higher seroprevalence of EBOV antibodies in *E. helvum* from Cameroon, with 107 out of 817 positive samples (*29*). At the time of writing this manuscript, the reservoir host of EBOV is yet to be determined, though bats have been heavily implicated (*30*).

Our study detected seroprevalence of neutralising antibodies against several viruses, but we cannot determine whether these viruses had sustained a prolonged infection in the bats, since RNA extractions were not carried out. Nonetheless, this raises the question as to whether the immune system of *E. helvum* is able to clear these viruses quickly, and how long the neutralising antibodies persist after infection. Ultimately, ongoing research on the dynamics of immune responses in bats (*31–33*) suggests that sero-surveillance studies provide a reliable assessment of potential reservoirs for these viruses.

In summary, our serological screening of *E. helvum* sera obtained from bats has revealed the presence of antibodies against several highly pathogenic viruses with epidemic potential. The capture and preparation of these bats for human consumption suggests a potential for direct exposure to bat bodily fluids, thereby elevating the risk of cross-species transmission of the viruses identified in this study. Given that human settlements are encroaching into areas known to harbour large bat colonies, the risk of zoonotic spillover will continue to increase. Therefore, monitoring of bat viruses especially in the large bat populations in Sub-Saharan Africa is crucial in order to better understand the prevalence and transmission of these viruses and to mitigate the risk of potential spillover events.

## Author Contributions

DC, AB, ESG and NT conceptualized the study. VOA and CTS collected and processed serum samples. DC, MN, BA, KDC, JDR, MD and SS carried out PVNA and data processing and analysis. AH generated figures and with AB and ESG analyzed the processed data, and provided expert critical feedback. All authors contributed significantly to the article and approved the final submitted version.

## Acknowledgements

Ethical approval for the collection of bat surveillance sera was received from the Animal Ethics Committee and Research Ethics Committee of University of Pretoria, with certificate numbers V092-18 and REC097-18, respectively. Approval to sample bat populations (VDS/194/S.4/11/T/85) was obtained from the Director/Chief Veterinary Officer of Nigeria, Department of Veterinary and Pest Control Services, Federal Ministry of Agriculture and Pest Control Services, Abuja Nigeria. A.C.B. was part funded by the UK Department for the Environment, Food and Rural Affairs (Defra) and the devolved Scottish and Welsh governments under grants SE2213 and SV3006.

## Biographical Sketch

Dr Cantoni, currently at the MRC-Centre for Virus Research, University of Glasgow, is interested mostly in SARS-CoV-2 and neglected tropical viruses; from both a serosurveillance and molecular biology perspective.

